# Genomic characterization of a wild-like tomato accession from Arizona

**DOI:** 10.1101/2022.02.11.480156

**Authors:** Jacob Barnett, Gina Buonauro, April Kuipers, Manoj Sapkota, Esther van der Knaap, Hamid Razifard

**Affiliations:** Graduate Program in Organismic and Evolutionary Biology, University of Massachusetts Amherst, Amherst, Massachusetts, 01003, USA; Biology Department, University of Massachusetts Amherst, Amherst, Massachusetts, 01003, USA; Scottsdale, Arizona, USA; Institute of Plant Breeding, Genetics and Genomics, University of Georgia, Athens, GA 30602, USA; Department of Horticulture, University of Georgia, Athens, GA 30602, USA; Center for Applied Genetic Technology, University of Georgia, Athens, GA 30602, USA; School of Integrative Plant Science, Cornell University, Ithaca, New York, 14850, USA

**Keywords:** Crop relatives, plant domestication, genomics, migration, *Solanum lycopersicum*, phylogenetics, tomato

## Abstract

Tomato domestication history has been revealed to be a highly complex story. A major contributor to this complexity is an evolutionary intermediate group (*Solanum lycopersicum* var. *cerasiforme* (Alef.) Voss; SLC) between the cultivated tomato (*Solanum lycopersicum* var. *lycopersicum* L.; SLL) and its wild relative (*Solanum pimpinellifolium* L.; SP). SLC includes accessions with a broad spectrum of genomic and phenotypic characteristics. Some of the SLC accessions were previously hypothesized to be spreading northward from South America into Mesoamerica and that migration probably entailed reversal to wild-like phenotypes such as smaller fruits. Prior to this study, the northernmost confirmed extension of the SLC was limited to northern Mexico.

In this study, we employed genomic methods to investigate the origin of a wild-like tomato found in a garden in Scottsdale Arizona, USA. The so-called “Arizona tomato” featured a vigorous growth habit and carried small fruits weighing 2-3 grams. Our phylogenomic analyses revealed the identity of the Arizona tomato as a member of the Mexican SLC population (SLC MEX). To our knowledge, this is the first report of an SLC accession, confirmed using genomics, growing spontaneously in Arizona. This finding could have implications for conservation biology as well as agriculture.

---

Crop relatives are an important reservoir of genetic diversity and are valuable sources of natural variation for crop improvement as well as insight into evolutionary processes (Turner-Hissong et al. 2020). Some semi-domesticated plants can also manifest wild-like phenotypes when growing in natural habitats and released from pressures of artificial selection, i.e., feralization. Feral crop relatives can sometimes possess weedy or invasive characteristics and could pose a threat to crops or native plant species outside of native habitats (Wu et al. 2021). Thus, the occurrence and spread of crop relatives is of interest to both agriculture and conservation of wild biodiversity, especially given the environmental shifts caused by climate change (IPCC 2021).

Tomato (*Solanum lycopersicum* L. var. *lycopersicum*, hereafter “SLL”) is one of the most valuable vegetable crops worldwide (FAOSTAT). Currently, wild and semi-wild tomato populations are found throughout western South America, Central America, and Mexico. *Solanum pimpinellifolium* L. (hereafter “SP”) is the closest wild relative of the common cultivated tomato and has been found in coastal to mid-elevation regions of Ecuador and Peru (Moyle 2008). *Solanum lycopersicum* L. var. *cerasiforme* (hereafter “SLC”) is an evolutionarily intermediate group between SP and SLL with an extensive geographic distribution from South America to Mexico, as well as diverse morphological and genomic features (Razifard et al. 2020). Colloquially, any tomato plants with fruits of intermediate size between wild and cultivated accessions are often called “cherry tomatoes”. However, the fruit size alone cannot reliably distinguish different tomato groups, considering that it is a highly variable trait depending on the environment (Lewis 2016).

Potentially wild-like “cherry tomatoes” have been found growing as weeds in California and Hawaii (R. Chetelat pers. comm.) and some of these plants have been collected as accessions in the C. M. Rick Tomato Genetics Resource Center at the University of California, Davis, USA (TGRC, http://tgrc.ucdavis.edu). Also, as of June 2022, there are a few records of “cherry tomato” from the continental US, listed on GBIF and iNaturalist (https://www.gbif.org/species/2930165 and https://www.inaturalist.org/observations?taxon_id=242951). Unfortunately, many of these collections do not contain sufficient morphological evidence for an accurate identification. However, one such collection from North Ridgeville, Ohio, represents a spontaneous plant resembling the commercial “cherry tomato”, and appears to have escaped from a garden or a tomato farm nearby. Importantly, none of these plants have been characterized genetically and thus their ancestry remains unclear.

Recent studies on the domestication history revealed that tomato has undergone a complex and dynamic domestication process likely involving initial domestication in South America, followed by northward spread of SLC to Mesoamerica (probably also involving feralization) and redomestication in Mexico (Blanca et al. 2015, Razifard et al. 2020). Other hypotheses suggested the northward spread of the SLC followed by the southward spread of some accessions carrying derived alleles for some of the domestication genes (Pereira et al. 2021, Blanca et al. 2022). Prior to this study, the northernmost confirmed extension of the SLC was limited to northern Mexico.

In this study, we employed genomic methods to investigate the origin of a wild-like tomato found growing spontaneously in a garden in Scottsdale Arizona, USA, arriving at an identification based on a combination of morphological and genomic evidence. We conclude by discussing the potential implications of this finding for conservation biology as well as agriculture.

## Materials and Methods

This study was motivated by an inquiry from April Kuipers, regarding an unusual tomato plant with wild-like phenotypes that was growing with minimal care in the yard of their house in Arizona (Arizona tomato). According to Ms. Kuipers, the tomato plant was not planted intentionally and was unlikely to have come from any previous gardening activity, as the yard had been covered in landscaping rock for at least the previous ten years. The plant had established itself with little care in the juvenile stage, although it was irrigated later along with other ornamental plants growing in the yard. Over time, the plant had grown into a relatively large bush entangled with a *Bougainvillea* and carried hundreds of small red fruits typical of those produced by SP (Fig. 1). Based on the wild-like phenotypes of small fruits (2-3 grams, ~1.7 cm diameter) with two locules, and the indeterminate growth habit, we hypothesized that the plant was either a wild (SP) or a wild-like tomato relative (SLC). It is often difficult to separate wild-like SLCs from SP based solely on morphology, considering the high level of resemblance between SP and wild-like SLC tomatoes. Therefore, we obtained genome sequence data from the “Arizona tomato” to arrive at a more reliable identification using both morphological and genomic evidence.

**Fig. 1.**
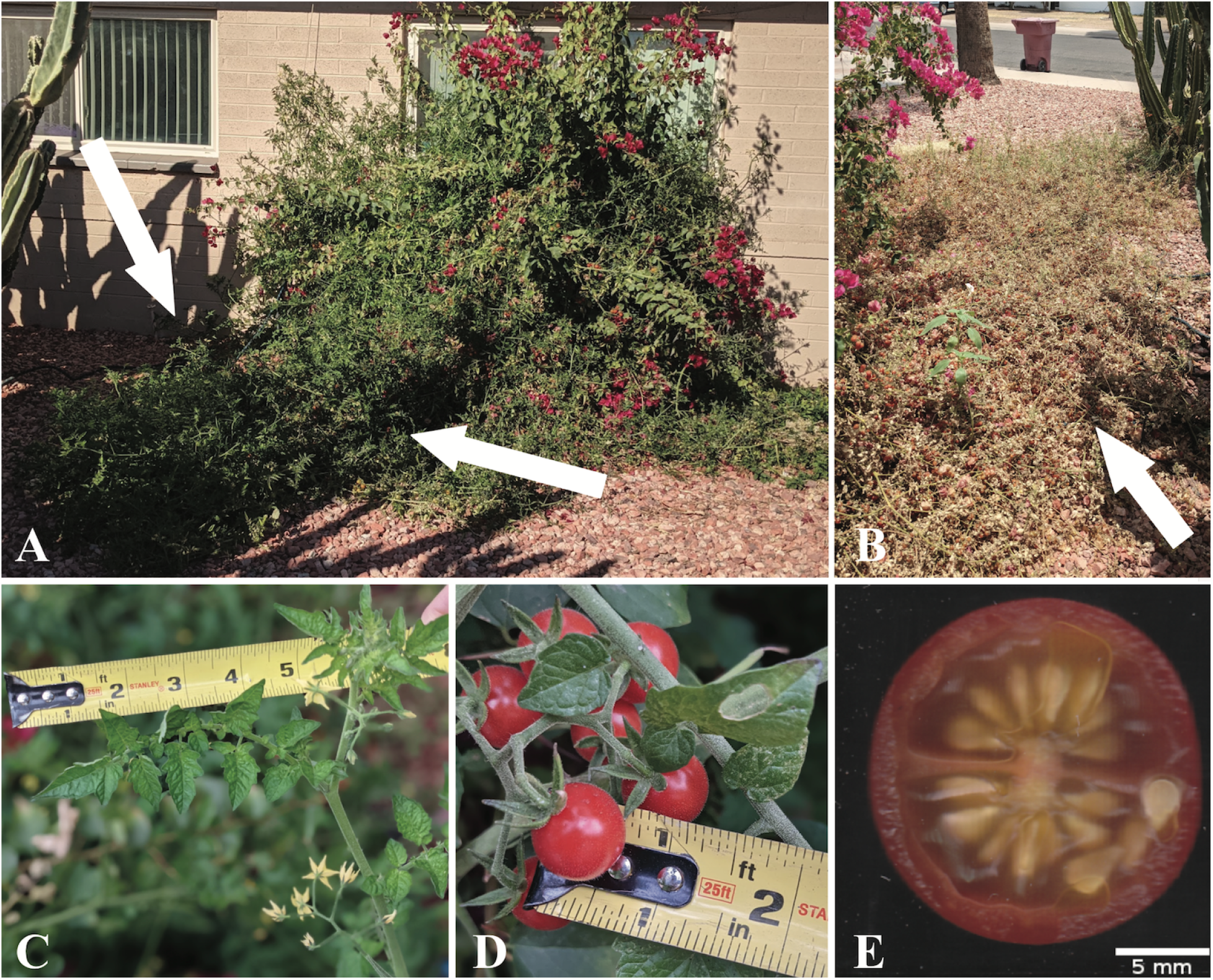
Plant at collection site in Scottsdale, Arizona, USA: A) Growing up and around a Bougainvillea bush with the majority of the plant visible in the lower left of the photo as indicated by white arrows, January 2021, and B) Extending outwards along ground as year progressed (indicated by white arrow), June 2021. C) Leaf and flowers in Arizona, January 2021. D) Ripe fruits in Arizona, January 2021. E) Scanned cross-section of ripe fruit grown at University of Massachusetts greenhouse, June 2021. Photos A-D by A. Kuipers, E by J. Barnett.

### Plant collection and voucher specimens

The “Arizona tomato” was discovered growing naturally in a suburban yard in Scottsdale, Arizona, USA in January 2021 by April Kuipers. All plant materials used in this study were obtained with permission from Ms. Kuipers, who found the plant on her personal property. She collected seeds from the fruits and sent them to the University of Massachusetts Amherst, where plants were grown in a greenhouse kept at 22 °C (with a few degrees of day/night fluctuations) under ambient daylight conditions from February - August 2021. Seeds were soaked in 2% sodium hypochlorite bleach solution, rinsed with distilled water, then placed ~ 0.5 cm deep in ProMix BX potting medium, resulting in 100% germination rate (nine out of nine seeds). Two plants were grown to maturity, and two specimens from each plant were preserved on June 30, 2021, and deposited in the University of Massachusetts Herbarium (MASS) as vouchers.

### Genome sequencing

Genomic DNA was extracted from a plant grown at Cornell University. Approximately 4 grams of leaf samples from the youngest leaves were used. Leaf samples were frozen and ground in liquid nitrogen. The DNA purification was conducted using NucleoBond® HMW DNA kit (from TaKaRa) and the purification steps were followed exactly as described in the protocol accompanying the kit. The total DNA yield from the leaf samples was 500 ng per microliter, of which 600 ng was used as input for DNA sequencing library preparation. The sequencing library preparation was conducted using New England Biolab FS DNA Library Prep Kit (E7805, E6177). A paired-end library was made with fragment sizes of 100-300 bp. The resulting library was visualized on a 0.4% agarose gel to confirm the expected spectrum of fragment sizes. The library was sequenced with Illumina NextSeq technology using the paired-end mode (2 × 150 bp) at the Genomics Facility of the Cornell Institute of Biotechnology.

### Sequence read alignment and variant calling

The reads obtained through genome sequencing were filtered by removing low-quality reads and the universal Illumina adaptors using cutadapt (v. 2.1; Martin 2011) with the default options. Remaining reads were checked for quality with FastQC (v. 0.11.8; https://www.bioinformatics.babraham.ac.uk/projects/fastqc/) and aligned to the most recent version of the cultivated tomato reference genome (SL4.0) using *mem* from bwa (v. 0.7.17; Li 2013). Similarly, genome alignments of SP, SLC, and SLL accessions created in previous studies (Razifard et al. 2020 and Pereira et al. 2021) were also included to create a variants dataset. A combined dataset of 21,358,376 variants including SNPs and indels (insertions and deletions < 10 bp) were made using *mpileup* from bcftools (v. 1.14; Danecek et al. 2021) software package. From this dataset, rare alleles (minimum allele frequency < 0.02) were removed using vcftools (v. 0.1.17; Danecek et al. 2011).

### Phylogenetic reconstruction

A SNP phylogeny was built using only four-fold degenerate SNPs (“4D SNPs”), i.e. SNPs at the third codon positions in which changes to all four base-pairs do not affect the translated amino acid from those codons. Such codon positions are considered to be under less selective pressure, thus more suitable for studying phylogenetic relationships. The input dataset for the phylogenetic analyses were made as follows: A custom script (available from https://github.com/hrazif) was developed in R (v. 4.0.5; https://www.r-project.org/) to extract 4D SNPs from the variants dataset described above. From the resulting 4D SNPs dataset, we kept only those with no missing data and minimum allele frequency > 0.012 (to keep 4D SNPs with alternate alleles in at least two homozygous accessions, four heterozygous accessions, or some other genotype combination). These steps resulted in 41,787 4D SNPs. For drawing a phylogenetic tree using the 4D SNPs, we used the coalescent method of SVDQuartets (Chifman and Kubatko, 2014) included in PAUP (v. 4a168; Swofford DL. 2003). The “exhaustive” search option, i.e. including all possible quartets, was chosen for the SVDQuartets analysis.

As an alternative method, we also generated a phylogeny based on a genome-wide dissimilarity matrix calculated from 13,956,415 variants (all variants except those with minimum allele frequency < 0.012 and missing data > 10%) in all accessions using SNPRelate (v1.10.2; Zheng et.al.,2012) in R. We then used the resulting distance matrix to create the phylogeny, based on the default “complete linkage” method.

### Geographic distributions

A map emphasizing the native distributions of the populations as well as the location of the newly discovered “Arizona tomato” was created using the Maps package (v3.2.0; https://CRAN.R-project.org/package=maps) in R.

## Results

### Genome sequencing and phylogeny

The genome sequencing of the “Arizona tomato” resulted in 227 million paired reads (2× 150 bp), corresponding to, on average, 41X coverage of the tomato genome. To create the input dataset for a phylogenetic reconstruction, we aligned the Arizona tomato genome sequences as well as the sequences created in Razifard et al. (2020) to the newest version of the tomato genome (SL4.0). Variant calling on the new genome as well as the genome sequences of SP, SLC, and SLL from Razifard et al. (2020) resulted in 21,358,376 SNPs, of which 13,956,415 variants were kept after excluding rare alleles.

We estimated the phylogeny of the accessions included in this study, using 41,787 SNPs in four-fold degenerate sites (Fig. 2). This phylogenetic reconstruction was consistent with the previously-published phylogenetic relationships between cultivated tomato and its wild and semi-wild relatives (Razifard et al. 2020). Briefly, we distinguished three main SP populations, from southern Ecuador (SP SECU), Peru (SP PER), and northern Ecuador (SP NECU). In Ecuador, there is genetic separation between the northern and southern SP populations (Fig. 2), with the southern Ecuadorian SP (SP SECU) being genetically closer to the Peruvian SP (SP PER) than to the SP population from northern Ecuador (SP NECU). The closest wild relative of SLC is SP NECU.

**Fig. 2.**
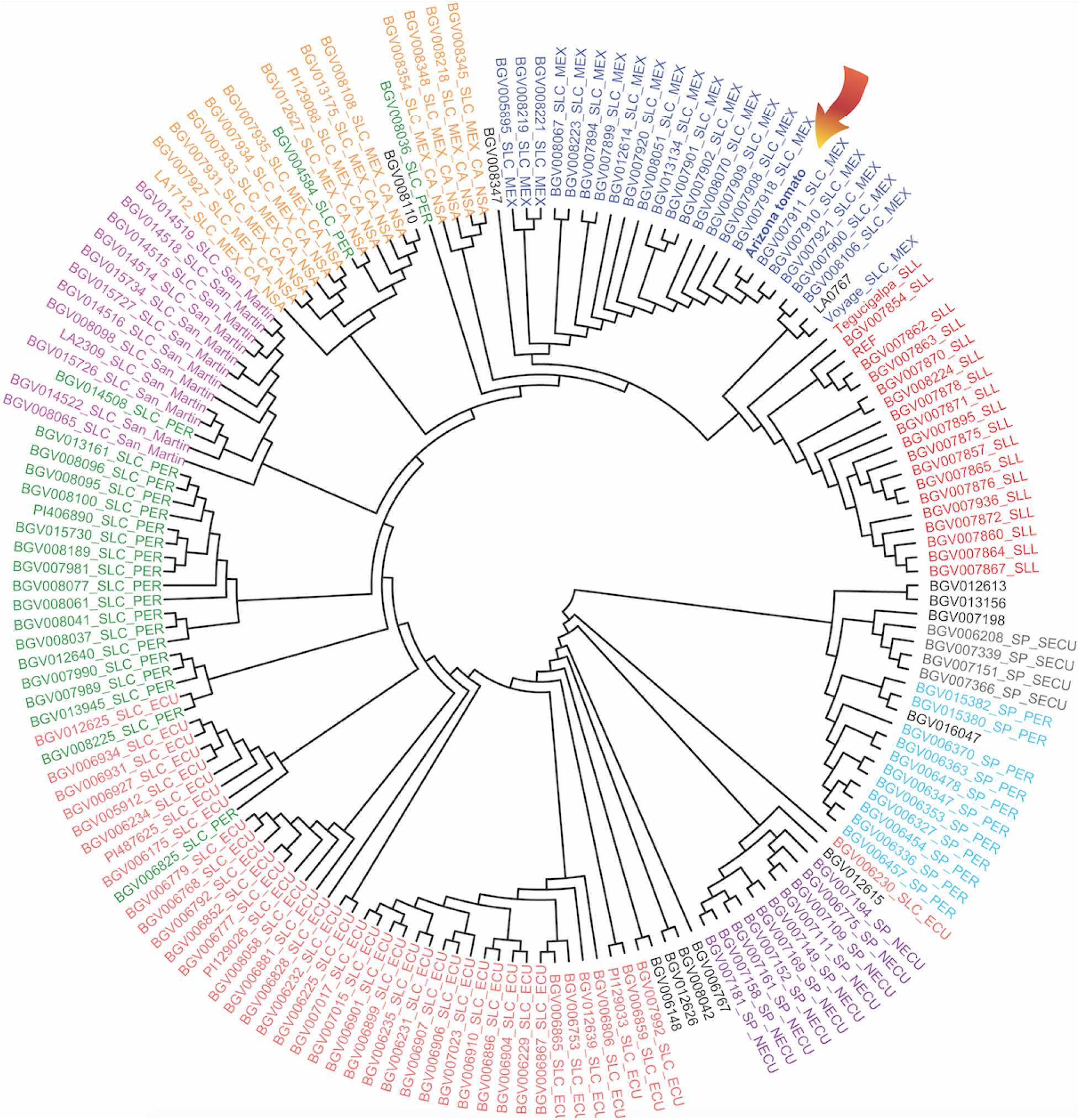
A reconstruction of the phylogenetic relationships between the cultivated tomato (*Solanum lycopersicum* var. *lycopersicum*, SLL) and its wild (*S. pimpinellifolium*, SP) and semi-wild relatives (*S. lycopersicum* var. *cerasiforme*, SLC). The phylogenetic position of the “Arizona tomato” is shown by an arrow. The definition of different populations of SP and SLC are according to Razifard et al. 2020, and colors are used to distinguish each population. Accessions without population assignments are those previously considered admixed accessions.

We also identified five main SLC populations, from Ecuador (SLC ECU), Peru (SLC PER), San Martin, Peru (SLC San Martin), an SLC population with a wide distribution range from northern South America to Mesoamerica (SLC MEX-CA-NSA), and the Mexican SLC population (SLC MEX). Two SLC populations, SLC MEX-CA-NSA and SLC MEX, are the northernmost extensions of SLC known to date.

The phylogenetic reconstruction revealed that the “Arizona tomato” was nested within a clade of SLC MEX accessions, indicating that it was genetically close to SLC accessions from Mexico. An alternative phylogenetic reconstruction based on genetic dissimilarity, using all ~ 14 million SNPs (Fig. 3), revealed the same position for the “Arizona tomato” - within SLC MEX.

**Fig. 3.**
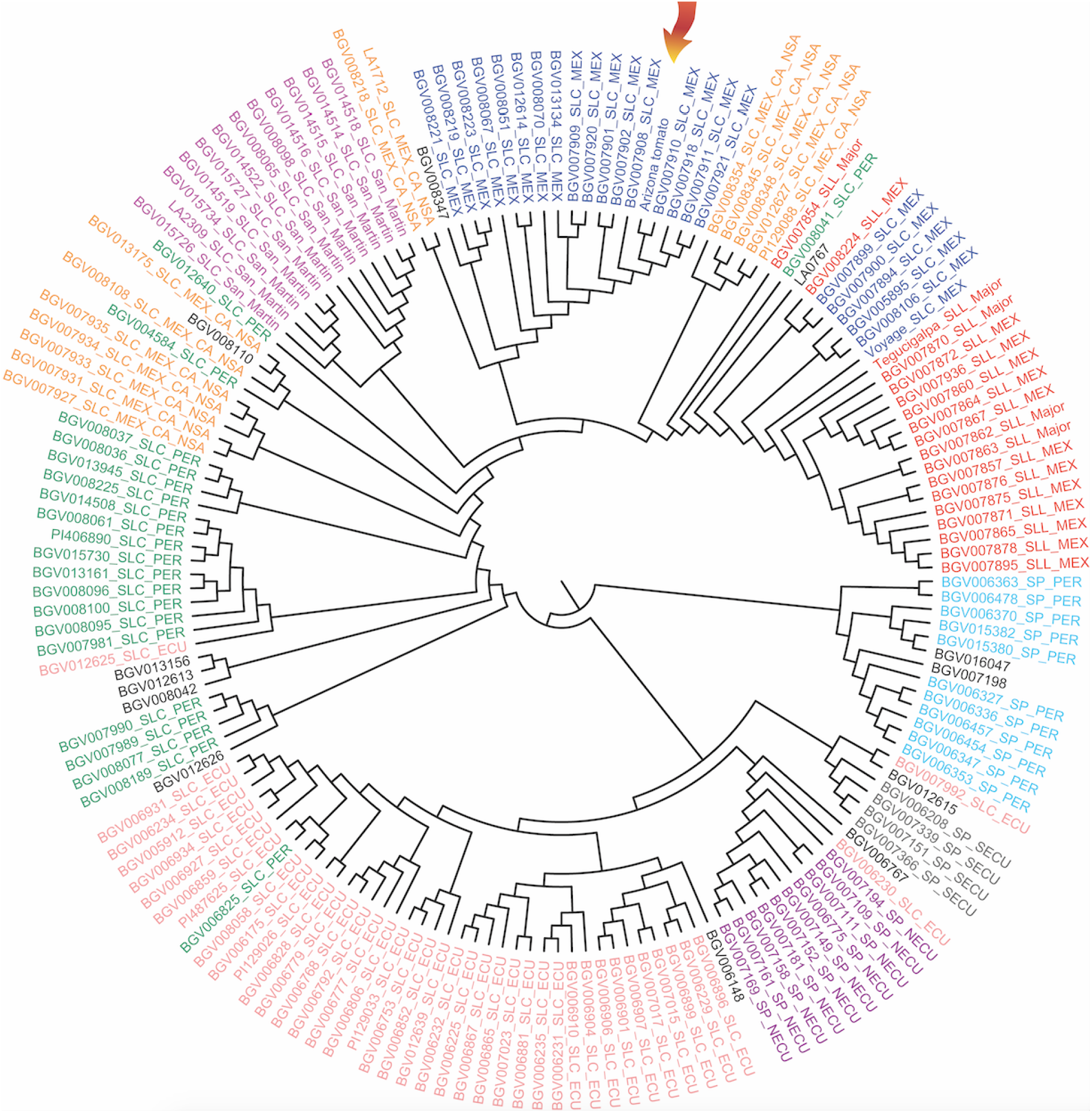
A distance-based reconstruction of the phylogenetic relationships between the cultivated tomato (*Solanum lycopersicum* var. *lycopersicum*, SLL) and its wild (*S. pimpinellifolium*, SP) and semi-wild relatives (*S. lycopersicum* var. *cerasiforme*, SLC). The phylogenetic position of the “Arizona tomato” is shown by an arrow. The definition of different populations of SP and SLC are according to Razifard et al. 2020, and colors are used to distinguish each population. Accessions without population assignments are those previously considered admixed accessions.

### Geographic distributions

Among the main groups included in this study, SLC appeared to have the broadest geographic range, extending from the southernmost regions of South America to Central America and Mexico (Fig. 4). The “Arizona tomato” reported in this study was genetically close to SLC MEX accessions, distributed in Eastern Mexico, even though it is geographically closer to accessions of SLC MEX-CA-NSA distributed in western Mexico.

**Fig. 4.**
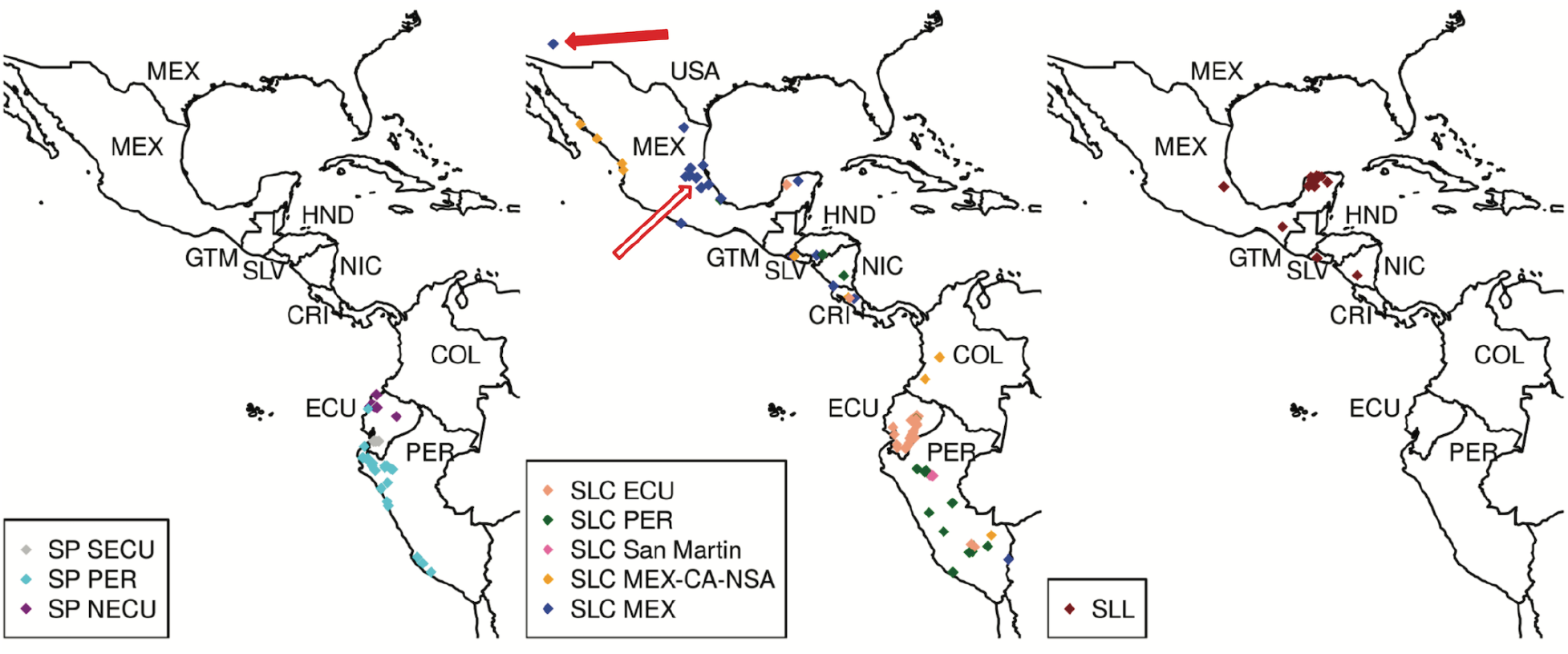
Distribution map of populations of wild (SP), semi-wild (SLC), and cultivated (SLL) tomato groups. Solid red arrow indicates collection location of the Arizona sample; outlined red arrow shows location of its putative sister population as identified by this study. Colors of points correspond to the population assignments shown in Figs. 2 and 3.

## Discussion

In this study, we report the results of phylogenomic and morphological analyses of the “Arizona tomato”, a tomato plant occurring spontaneously in the southern United States. Our studies revealed the identity of the “Arizona tomato” as a Mexican SLC. To our knowledge this represents the first detailed genomic characterization of a tomato relative in the United States, expanding the known northern range of SLC beyond Mexico.

Morphologically, wild-like SLCs can be difficult to distinguish from SP, the wild tomato relative. Having small fruits (2-3 grams, ~1.7 cm diameter) with two carpels, lobed leaflets as well as indeterminate growth habits (Fig. 1), the “Arizona tomato” resembled accessions of wild SP and wild-like SLC MEX-CA-NSA and SLC MEX (Razifard et al. 2020). Using genome sequencing, we reliably identified the ‘Arizona tomato” as a member of the Mexican SLC population (SLC MEX) found in eastern Mexico (Fig. 4). Previous studies identified the SLC MEX as the closest wild relative of the common cultivated tomato (SLL; Razifard et al. 2020). Two SLC populations, SLC MEX-CA-NSA and SLC MEX, are the northernmost extensions of SLC. It has been hypothesized that SLC arose approximately 78,000 years ago with northward spreads approximately 13,000 years ago (Razifard et al. 2020).

The confirmation of SLC MEX in the USA could represent a northward range expansion of this population by an as-yet unknown vector. Birds may have been an agent of seed dispersal, as the red fruits are an attractive food source for birds and are small enough to be consumed and potentially carried widely while being digested. For example, a previous study (Heleno et al. 2011) indicated that the wild tomatoes endemic to the Galapagos were probably first carried to the islands from the mainland by birds. According to Ms. Kuipers’ observations of the “Arizona tomato”, birds displayed an interest in the fruits. Arizona and Mexico lie along the migratory route of many bird species that could potentially carry seeds of some plant species (La Sorte et al. 2016).

Humans were also a plausible vector of spread, as several varieties of “wild” tomatoes are sold online to gardeners in present times. Personal observations from farmers’ markets in New York and Massachusetts suggest an interest in growing wild or wild-like tomatoes as a hobby and/or a small business opportunity in the US, as their desirable fruit flavor, small fruit size, and vigorous fruit production make for attractive selling points. One of the commercially available varieties known as “Matt’s Wild Cherry” (Johnny’s Selected Seeds) reportedly originated from Hidalgo, Mexico, which lies within the known range of SLC MEX. Another variety called “Texas Wild” (TomatoFest®) is claimed to have been collected from a patch of “wild” tomatoes in southern Texas. “Red Currant Tomato” seeds labeled as wild SP are also available (Victory Seeds®). Further phylogenomic work is needed to confirm the genetic identity of these seeds, but their sale could be at least partially responsible for the spread of wild or wild-like tomatoes into new areas.

Regardless of the exact manner of its spread, the spontaneous appearance of the “Arizona tomato” could have implications for both agriculture and conservation biology. In Mexico, accessions of SLC MEX and SLC MEX-CA-NSA are often found in association with houses and farms (Razifard et al. 2020). Occasionally, these plants appear in fields of crops such as maize and might be foraged, considering their edible fruit (Casas et al. 2016). In some other cases, however, feral or de-domesticated crops have become threats to crops and native species due to their weedy traits (Wu et al. 2021). While the environmental and agricultural risk of this new introduction is currently unknown, wild-like tomatoes could potentially still compete with native plants if they became established in natural habitats. This is an important possibility to consider given the shifting ranges of many plants due to climate change (Pecl et al. 2017).

## Conclusion

In the face of a changing climate, it is likely for plant species to spread to previously uninhabitable environments. Although some such spreads may have a short-term economical benefit, they may cause long-lasting harm to the native ecosystems as well as agriculture. In this study, we for the first time confirmed the introduction of a Mexican SLC tomato to the US. Although the impact of this introduction remains to be studied, we recognize the need to balance the preservation of crop relative germplasm with the potential risks of establishment in new territories.

## Acknowledgments

We thank Margaret H. Frank (Cornell University) for kindly providing reagents for genome library preparation. This research was funded by NSF IOS 1942437 to Margaret Frank and NSF IOS 1564366 to JB, MS and EV. We also thank Roberta Lombardi at the University of Massachusetts Herbarium (MASS) for assisting with preparation and deposition of the voucher material.

## Author Contributions

AK discovered the plant, provided photographs, and sent seeds to JB and HR. JB and GB grew plants from seed and prepared the voucher specimens. HR extracted and sequenced DNA and performed the genomic analyses. MS and EV aligned the previously sequenced genomes to the newest version of the tomato reference genome. JB and HR wrote the manuscript. GB, AK, MS, and EV provided feedback on the manuscript.

## Appendix 1.

Voucher information for the specimens that produced phenotyped fruits, and NCBI Sequence Read Archive (SRA) information for the specimen used in molecular phylogenetic analyses. Description strings are organized as follows: taxon, herbarium voucher, herbarium institution code, NCBI SRA BioProject ID. Voucher specimens are housed at the herbarium of the University of Massachusetts Natural History Collections (MASS).

*Solanum lycopersicum* var. *cerasiforme*, Plant # 1 001 barcode 00437024, MASS, —; *Solanum lycopersicum* var. *cerasiforme*, Plant # 1 002 barcode 00437025, MASS, —; *Solanum lycopersicum* var. *cerasiforme*, Plant # 2 001 barcode 00437022, MASS, —; *Solanum lycopersicum* var. *cerasiforme*, Plant # 2 002 barcode 00437023, MASS, —; *Solanum lycopersicum* var. *cerasiforme*, —, —, PRJNA802260.

